# Binary classification machine-learning Unveils Sex-Dependent mutated gene Signatures in Melanoma

**DOI:** 10.1101/2023.04.04.535515

**Authors:** Nadav Elkoshi, Shivang Parikh, Sajeda Mahameed, Abraham Meidan, Eitan Rubin, Carmit Levy

## Abstract

There are significant differences in the prevalence of cancer type, primary tumor body site, and mutation load between men and women, but the mechanisms underlying these sex-dependent differences is mostly unknown. Here we used binary classification machine-learning methodology to study sex-correlated somatic mutations signatures in cutaneous melanoma. We identified a number of genes that are more frequently mutated in females compared to males. Mutations in two genes, *LAMA2* and *TPTE*, together with a set of specific genes that are not mutated, can predict sex of melanoma patients. Over representation analysis of genes clustered with *LAMA2* revealed significant enrichment in androgen and estrogen biosynthesis and metabolism pathways, suggesting that mutation of *LAMA2* might be involved in biased sex hormone synthesis in melanoma. Taken together, our analysis shows that gender can be predicted based on mutation status of genes in melanoma and that certain mutations are predictive of survival beyond sex differences. Our results will lead to better diagnosis and more effective personalized treatment of melanoma.

**Summary:** It was observed that between men and women there is a significant difference in the prevalence of cancer type, the primary tumor body site, and the number of mutations found in a given tumor type. However, the mechanisms behind these gender differences are mostly unknown. To investigate sex corelated somatic mutation signatures in cutaneous melanoma we used binary classification machine-learning methodology. We identified specific genes that are more frequently mutated in females compared to males, including *LAMA2* and *TPTE*, which are predictive of gender. We also found a significant enrichment in androgen and estrogen biosynthesis and metabolism pathways clustered with *LAMA2*, suggesting that mutation of *LAMA2* might be involved in biased sex hormone synthesis in melanoma. We showed that gender can be predicted based on mutation status of genes in melanoma and that certain mutations are predictive of survival. Our findings could lead to better diagnosis and more effective personalized treatment of melanoma.

## Introduction

Sex-specific differences are observed in the mortality and the incidence of cancer [1]. Males have overall higher incidence and mortality compared to females [2]. Males have higher occurrence of colorectal [3], stomach [4], liver [5], and bladder [6] cancers and leukemia [7], whereas females have increased occurrence of thyroid cancer [8]. Primary cancer body location is also gender biased in melanoma [9] and colorectal cancer [10]. Analysis of the body distribution of primary cutaneous melanoma showed that men are more likely to have tumors in the trunk regions, especially on the back, whereas women have more tumors in the lower body [11]. Proximal, compared to distal colon or rectal, cancer rates were higher among females than males for young blacks compared to other races [10]. The causes and the mechanisms behind cancer phenotype disparities between males and females remain to be largely unknown.

Further, gender disparities in mutation load were observed in both lung cancer [12] and melanoma [13]. Among lung adenocarcinoma patients who do not smoke, males have significantly more mutations than females [12], whereas melanoma tumors in males harbor significantly more missense mutations compared to tumors in females [13]. Malignant melanoma is the most aggressive form of skin cancer and accounts for approximately 325,000 cases estimated globally as per the statistics of the year 2020 [14]. UV radiation is a known driver of cutaneous melanoma [15], and UV mutational signatures characteristic of this malignancy have been identified [16,17]. Although it has been suggested that sex disparities in melanoma arise from different behavioral features such as sun-seeking behavior [18,19], clothing [20], and skin screening [20], some intrinsic biological factors have been identified [21–23]. Further, elevated mutation burden in melanoma is strongly associated with better survival [24] and with increased effect in females compared to males [13], demonstrating sex-related differences in disease formation and development. Although acknowledged as crucial in clinical medicine, the impact of sex on cancer susceptibility, progression, therapeutic response, and survival remains poorly researched [2]. Therefore, a better understanding of sex disparity in cancer initiation, development, and progression is required to advance in cancer prevention, treatment, and outcome.

In this study, we investigated whether there is a sex disparity in somatic mutations in melanoma using the machine-learning-based software WizWhy for binary classification [25]. We identified genes that are more frequently mutated in females than in males, from which only two, *LAMA2* and *TPTE*, when appear together with a set of specific genes in their non-mutated statues, are predictive of sex. Over representation analysis of *LAMA2*’s cluster genes revealed 3 enriched pathways, in which androgen and estrogen biosynthesis and metabolism was the most significant, suggesting that mutated *LAMA2* might be involved with gender biased sex hormones synthesis in melanoma. Our analysis demonstrated that mutation status in melanoma depends on gender and that mutation status is predictive of survival. These results further our understanding of differences between men and women in mutational burden and melanoma incidence and prognosis.

## Methods

### Data and pre-processing

For analysis, we downloaded the TCGA melanoma somatic mutations data from the NIH GDC portal, accessed at https://portal.gdc.cancer.gov/. Synonymous and rare mutations (occurring in <10% of males and females) were discarded. Single-nucleotide polymorphism data was projected into gene space and binarized, so that each gene was considered mutated in an individual if it contained one or more mutations after filtering. The number of mutated genes per individual was binned (with each bin including 7 sequential counts), and individuals were randomly paired by their gender (one male, one female) and mutated gene count bins. Pairs were trimmed until no differences in clinical and demographic features (Supplementary Table 1) were detected using the appropriate test (with multiple testing correction for variables with multiple levels). Finally, a paired *t*-test was also used to test for differences in raw total mutated gene counts, and no significant difference was detected.

### Machine learning and evaluation

Pairs were randomly assigned to training (70%) and testing (30%) datasets four times. Each dataset was used to create a predictive model using the WizWhy binary classification software [25]. Briefly, the algorithm identifies if-and-only-if and if-then predictive rules, specifying its answer as rule sets. A case is assigned to one class if any of the rules of the set are true; if none of the rules are true, the case is assigned to the other class. In this study, only if-then predictive rules were detected.

Machine learning code will be available upon request.

### Statistics and implementation

Unless otherwise specified, a p value cutoff of ≤ 0.05 was considered significant. The Benjamini-Hochberg false discovery rate correction [26] was used for multiple testing correction unless otherwise specified. The number of mutated genes, age at diagnosis, and Breslow thickness at diagnosis were assessed using a paired *t*-test, and other features (ethnicity, race, tumor status, vital status, AJCC stage, node involvement, metastasis, tumor site, primary location, Clark level at diagnosis, tissue source, and ulceration) were assessed using McNemar symmetry tests for paired contingency tables with adjusting for multiple p values with the nominalSymmetryTest from the rcompanion package in R. Paired data with one or more missing values were removed before applying the statistical tests. Analyses were done in R, except for the learning step, which was conducted with a special version of the WizWhy software that was adjusted to use many features by storing some intermediates on disk instead of in memory. Over representation analysis and induced network analysis were performed using the ConsensusPathDB [27] available at http://consensuspathdb.org.

## Results

A careful pre-processed of TCGA mutation data yielded 159 males and 159 females paired according to the number of mutated genes, ensuring that pair-wise data were not statistically biased over multiple clinical and demographic features (Supplementary Table 1). Pairs were randomly split four times for training (70% of the male-female pairs) and testing sets (30% of the male-female pairs), and the training set was used to train a predictive model using the WizWhy algorithm. Using the test sets, we estimated the accuracy of the model to be 75.2% ± 4.2% (mean ± standard deviation). This indicated that sex could be predicted from somatic non-synonymous mutations in melanoma tumors (Fig. 1a).

**Figure 1:**
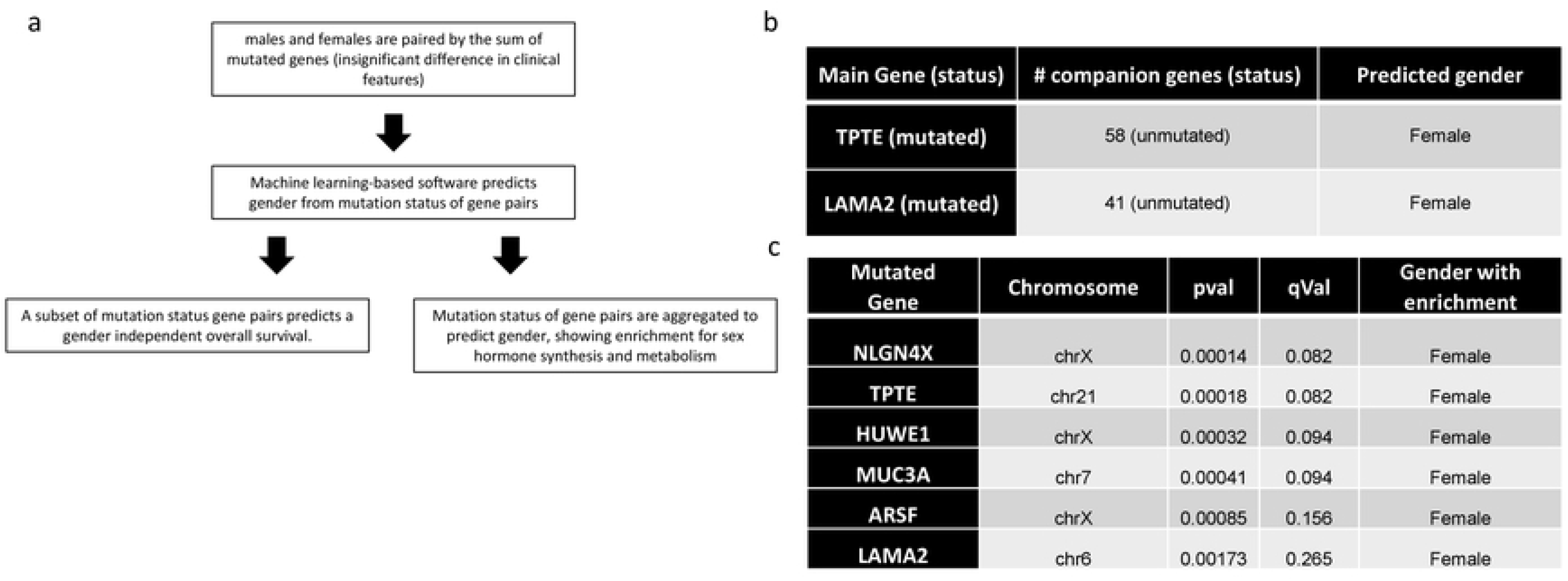
machine learning algorithm identifies gender patients by the occurrence of LAMA2 mutations paired with 41 unmutated genes. a) Workflow schematics. b) Patients’ gender is predicted to be female from mutation status of gene pairs. Gene pairs aggregated into common gender segregators are depicted (n=4). c) Top 3 genes harboring a gender dependent mutation status bias in melanoma.

To identify mutation patterns that can robustly predict gender, unstable predictive rules that did not repeat in all four random subsamples were discarded, and genes were clustered transitively according to predictive laws by common gene mutation status and prediction outcome. For example, the rules a) “if gene A is mutated and gene B is not mutated then gender is female” and b) “if gene A is mutated and gene C is not mutated then gender is female” can be clustered into the common rule c) “if gene A is mutated and gene B or gene C are not mutated, then gender is female”. Using this approach, rules could be grouped into two clusters, one with *TPTE* as the main hub, and one with *LAMA2* as the main hub (Fig. 1b). *TPTE* occurred in 58 stable rules and *LAMA2* in 41 stable rules. It should be noted that all stable rules contained either *TPTE* or *LAMA2*.

When the genes were ranked by their differences between males and females *TPTE* was ranked second, and *LAMA2* was ranked sixth; the p value for *LAMA2* was approximately 10-fold larger than that for *TPTE* (Fig. 1c and Supplementary Table 2). Therefore, we focused on rules involving *LAMA2* rather than *TPTE*.

*Laminin subunit alpha 2 (LAMA2*) encodes a laminin subunit and was previously suggested to be a tumor suppressor (Akhavan et al. 2012; Wang et al. 2019). *LAMA2* expression is downregulated due to promoter hypermethylation in invasive pituitary adenomas and colon and bladder cancers [28,29]. Further, LAMA2 modulates PTEN to affect the PI3K/AKT pathway [28] and it is a tumor suppressor in lung adenocarcinoma [30].

Over-representation analysis of genes clustered with *LAMA2* revealed three enriched pathways (with q values of 0.0879), in which androgen and estrogen biosynthesis and metabolism was the most significant. This suggests that mutated *LAMA2* might be involved with sex hormone synthesis, which might impact melanoma progression (Fig. 2a). Characterization of induced network modules of *LAMA2* and its clustered genes revealed that protein interactions (especially with some protein intermediates) were the most common connections in the network (Fig. 2b), suggesting that genes in this cluster may facilitate physical connections at the proteomic level.

**Figure 2.**
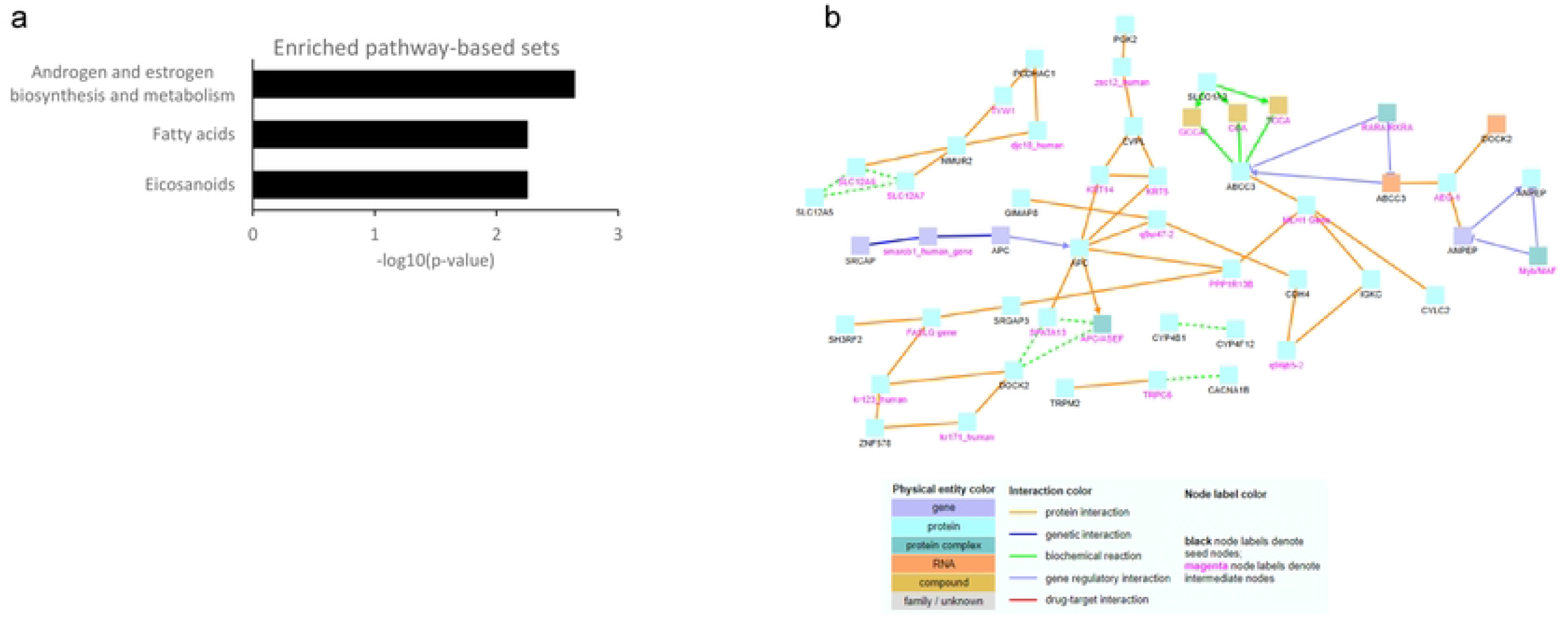
Enriched pathways of LAMA2 gender segregators and mutations status survival prediction. a) LAMA2 unmutated co-partners enriched pathways b) induced network of LAMA2 with its unmutated co-parteners

Next, we asked whether the gender segregation rules including *LAMA2* are predictive of survival. We tested each of the 41 rules for survival prediction using log-rank test following multiple test correction and found that among patients with *LAMA2* mutations, unmutated *TENM2* was predictive of significantly better overall survival compared to the mutated form (Fig. 3). No other rules were significant for overall survival prediction. We therefore conclude that our mutation-based gender segregation rules may play, to some extent, a role in melanoma survival beyond gender differences.

**Figure 3:**
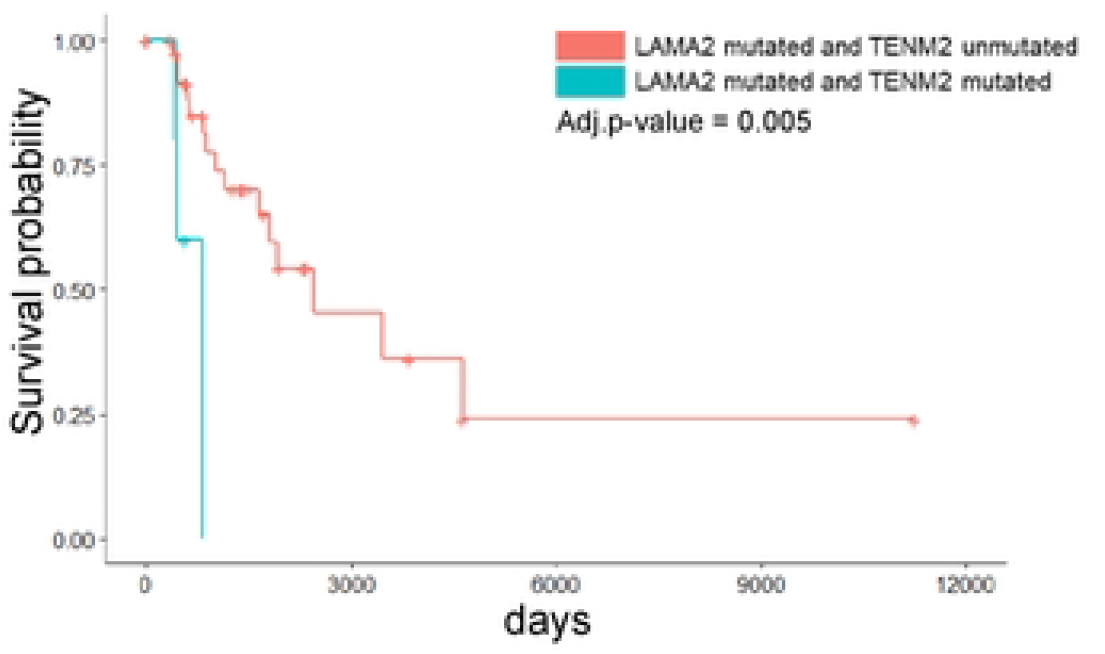
Paired mutations status predicts survival. TENM2 mutation status predicts survival of melanoma patients with mutations in LAMA2

## Notes

### Competing Interest Statement

The authors have declared no competing interest.

